# Triterpene RDF: Developing a database of plant enzymes and transcription factors involved in triterpene biosynthesis using the Resource Description Framework

**DOI:** 10.1101/2024.01.08.574260

**Authors:** Keita Tamura, Hirokazu Chiba, Hidemasa Bono

**Affiliations:** Genome Editing Innovation Center, Hiroshima University, Higashihiroshima, Hiroshima 739-0046, Japan; Database Center for Life Science, Joint Support-Center for Data Science Research, Research Organization of Information and Systems, Kashiwa, Chiba 277-0871, Japan; Graduate School of Integrated Sciences for Life, Hiroshima University, Higashihiroshima, Hiroshima 739-0046, Japan

**Keywords:** annotation, database, SPARQL, Resource Description Framework, triterpene

## Abstract

Plants produce structurally diverse triterpenes (triterpenoids and steroids). Their biosynthesis occurs from a common precursor, namely 2,3-oxidosqualene, followed by cyclization catalyzed by oxidosqualene cyclases (OSCs) to yield various triterpene skeletons. Steroids, which are biosynthesized from cycloartenol or lanosterol, are essential primary metabolites in most plant species, along with lineage-specific steroids, such as steroidal glycoalkaloids found in the *Solanum* species. Other diverse triterpene skeletons are converted into triterpenoids, often classified as specialized compounds that are biosynthesized only in a limited number of plant species with tissue-or cell-type-specific accumulation in plants. Recent studies have identified various tailoring enzymes involved in the structural diversification of triterpenes as well as transcription factors that regulate the expression of these enzymes. However, the coverage of these proteins is scarce in publicly available databases for curated proteins or enzymes, which complicates the functional annotation of newly assembled genomes or transcriptome sequences. Here, we created the Triterpene RDF, a manually curated database of enzymes and transcription factors involved in plant triterpene biosynthesis. The database (https://github.com/ktamura2021/triterpene_rdf/) contains 526 proteins, with links to the UniProt Knowledgebase or NCBI protein database, and it enables direct download of a set of protein sequences filtered by protein type or taxonomy. Triterpene RDF will enhance the functional annotation of enzymes and regulatory elements for triterpene biosynthesis, in a current expansion of availability of genomic information on various plant species.

## (Introduction)

Triterpenes are among the most diverse groups of natural compounds found in plants. These compounds are derived from six isoprene units. The last common triterpene precursor is 2,3-oxidosqualene, from which oxidosqualene cyclases (OSCs) generate more than 100 different triterpene skeletons (Sawai and Saito 2011; Thimmappa et al. 2014). Among the triterpene skeletons, cycloartenol and lanosterol are intermediates in the biosynthesis of phytosterols, which are indispensable components of plasma membranes or the plant hormone brassinosteroids (Ohyama et al. 2009, 2007). In addition to these primary metabolites, many specialized metabolites with beneficial bioactivities are known as triterpenes, including glycyrrhizin and ginsenosides, which are derived from β-amyrin and dammarenediol-II, respectively (Sawai and Saito 2011; Seki et al. 2015). Although the definition is not uniform, we classified triterpenes into steroids, which are compounds derived from cycloartenol or lanosterol, and triterpenoids, which are compounds derived from other triterpene skeletons, according to the classification by Ohyama et al. (2007).

Triterpene skeletons are further modified by several classes of enzymes to produce highly diverse structures of triterpenoids and steroids (Thimmappa et al. 2014; Sawai and Saito 2011). The biosynthetic pathways of phytosterols, including brassinosteroids, are relatively well understood, with the support of mutants of *Arabidopsis thaliana* (Bishop and Yokota 2001; Benveniste 2004). In contrast, enzymes for the modification of triterpene skeletons for the biosynthesis of triterpenoids remained poorly understood until the first identification of cytochrome P450 monooxygenase (P450) CYP93E1 in *Glycine max* (Shibuya et al. 2006) and UDP-dependent glycosyltransferases (UGTs) UGT73K1 and UGT71G1 in *Medicago truncatula* (Achnine et al. 2005). Since then, P450s and UGTs have been known to be central players in the structural diversification of triterpenoids, as P450s introduce functional groups, such as hydroxyl and carboxyl groups, into triterpene skeletons, and UGTs add sugar moieties to triterpene aglycones (Seki et al. 2015). In addition to these two protein families, recent studies have identified new types of enzymes involved in triterpenoid biosynthesis, including cellulose synthase-like glycosyltransferases (Chung et al. 2020; Jozwiak et al. 2020) and BAHD acetyltransferases (Kumar et al. 2021). Additionally, transcription factors (TFs) that regulate triterpenoid or steroid biosynthetic genes have been reported in a few biosynthetic pathways (Dinday and Ghosh 2023).

With the rapid progress in sequencing technologies, including long-read sequencing platforms, it is now possible to assemble large and complex plant genomes in more feasible ways (Sahu and Liu 2023). This has accelerated the elucidation of genome sequences of non-model plant species, including rare medicinal plants that produce valuable triterpenoids. In general, the functional annotation of predicted genes in assembled genomes after annotation of gene structures requires a set of well-curated reference proteome sequences, such as the reviewed (Swiss-Prot) part of the UniProt Knowledgebase (UniProtKB) (The UniProt Consortium 2023) and The Arabidopsis Information Resource (TAIR) (Berardini et al. 2015). Although these databases mainly cover OSCs or well-studied Arabidopsis biosynthetic pathways, most specialized triterpenoid biosynthetic pathway proteins have not been fully curated, which complicates functional annotation of triterpenoid biosynthetic genes in plant genomes. The TriForC database (http://bioinformatics.psb.ugent.be/triforc/ (Accessed Dec 21, 2023)) is a manually curated resource for enzymes involved in triterpene biosynthesis (Miettinen et al. 2018). Although this is a valuable resource to study plant triterpene biosynthesis, it does not directly provide a set of protein sequences from the database that would be useful for the functional annotation of predicted genes. The purpose of this study is to develop a database of functionally characterized proteins involved in triterpenoid and steroid biosynthesis in plants from public repositories, including UniProtKB, and to easily curate a set of protein sequences for the functional annotation of potential genes involved in triterpenoid or steroid biosynthesis.

## (Materials and methods)

The overall scheme of database construction is shown in Figure 1. By manual curation of the literature, with the help of previously published well-summarized review articles (Thimmappa et al. 2014; Malhotra and Franke 2022; Rahimi et al. 2019; Dinday and Ghosh 2023), we first created tables containing 526 proteins (v20231129_db02.tsv and v20231224_db02.tsv), indexed as TP0001–TP0526. These proteins were classified into five types: enzymes classified as OSC, P450, UGT, and “other enzyme”, and TF. Additionally, nine squalene cyclases (SCs) identified in ferns were added for reference (indicated as “*SC”). We classified these proteins according to their biosynthetic pathways. All proteins (except for SCs) were classified as either “triterpenoid” or “steroid” (forming the “pathway” column). Proteins classified as “triterpenoid” were further classified based on the triterpene skeleton compound produced by OSCs relating to the pathway (forming the “skeleton” column). We summarized the characterized function of the protein in the “function” column. In case of OSCs producing multiple triterpene skeletons, the characterization was made based on the major products. Each protein entry was linked to the accession of UniProtKB (forming the “uniport” column), using the accession or sequence indicated in the literature. Since UniProtKB distributes its databases in the Resource Description Framework (RDF) format, which allows the linking of various life science data using a standard query language (SPARQL) (The UniProt Consortium 2017; Jupp et al. 2014), we chose UniProtKB as the primary accession for each protein entry. For the entries that cannot map to UniProtKB accessions, NCBI protein database accessions were indicated (forming the “ncbiprotein” column). A primary citation for each entry was indicated as the PubMed accession (forming the “pubmed” column). Additional citations, Digital Object Identifiers for publications not available in the PubMed database, or notes for the entries were indicated in the “note” column.

**Figure 1.**
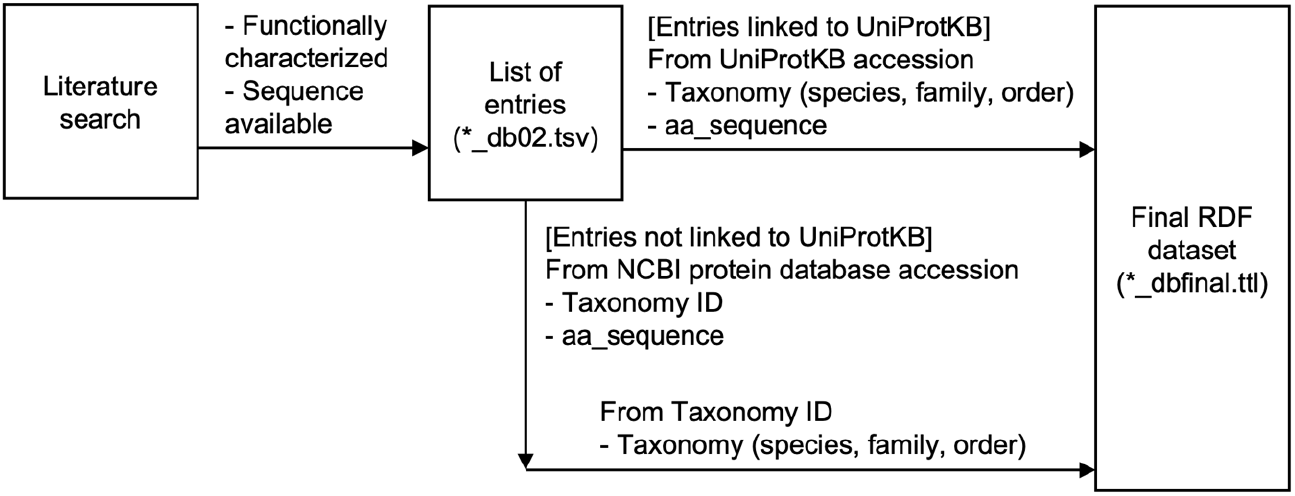
Overall scheme of the database construction.

Taxonomic and sequence information was retrieved using the accession of UniProtKB or NCBI protein database. For entries linked to UniProtKB, taxonomic information (scientific names of species, family, and order) and amino acid sequences (canonical isoforms) were retrieved using federated queries at the SPARQL endpoint of the UniProt database available at the RDF portal (https://rdfportal.org/sib/sparql; version 2023_02) (Kawashima et al. 2018). For entries linked to the NCBI protein database, taxonomy ID and amino acid sequences were retrieved by programmatic access using TogoWS (Katayama et al. 2010), and taxonomic information (scientific names of species, family, and order) was retrieved using the same method as entries linked to UniProtKB. The obtained taxonomic and sequence information was merged with the primary tables (v20231129_db02.tsv and v20231224_db02.tsv) to curate a final database table (v20231224_dbfinal.tsv) and a corresponding RDF dataset (v20231224_dbfinal.ttl), which we named Triterpene RDF. Triterpene RDF is accessible via a website (https://ktamura2021.github.io/triterpene_rdf/), which internally uses SPARQL query language against the RDF dataset. The sources of the database, custom scripts, SPARQL queries, and intermediate files for the construction of the database are available at the GitHub repository (https://github.com/ktamura2021/triterpene_rdf/).

## (Results and discussion)

A screenshot of the Triterpene RDF is shown in Figure 2. When users access a website, all entries are displayed first. Drop-down menus at the top of the page are provided to filter entries. The “Download FASTA” button enables users to download amino acid sequences for the displayed entries in FASTA format. Table 1 shows the number of protein entries in the database classified by “type” and “pathway”. We collected 397 and 120 entries for triterpenoid and steroid biosynthesis, respectively. P450s and OSCs were most commonly identified in both triterpenoid and steroid biosynthesis. The portion of entries labelled as “other_enzyme” in steroid biosynthesis is relatively higher, due to the involvement of many methyltransferases, reductases, and isomerases involved in phytosterol biosynthesis. We also analyzed the number of entries into triterpenoid and steroid biosynthesis, classified according to plant order (Tables 2 and 3). The most studied plant order for triterpenoid biosynthesis was Fabales (97 entries), followed by Brassicales (49 entries). All five proteins were identified in both orders (Table 2). The Fabales order includes the well-studied Fabaceae species for triterpenoid saponin biosynthesis, such as *Medicago truncatula* (25 entries) and *Glycine max* (15 entries). The most studied plant order for steroid biosynthesis was Brassicales (30 entries), followed by Solanales (27 entries) (Table 3). This reflects the elucidation of phytosterol biosynthetic pathways using *A. thaliana* and the recent accumulation of knowledge on steroidal glycoalkaloid biosynthesis in the *Solanum* species (Akiyama et al. 2023).

**Figure 2.**
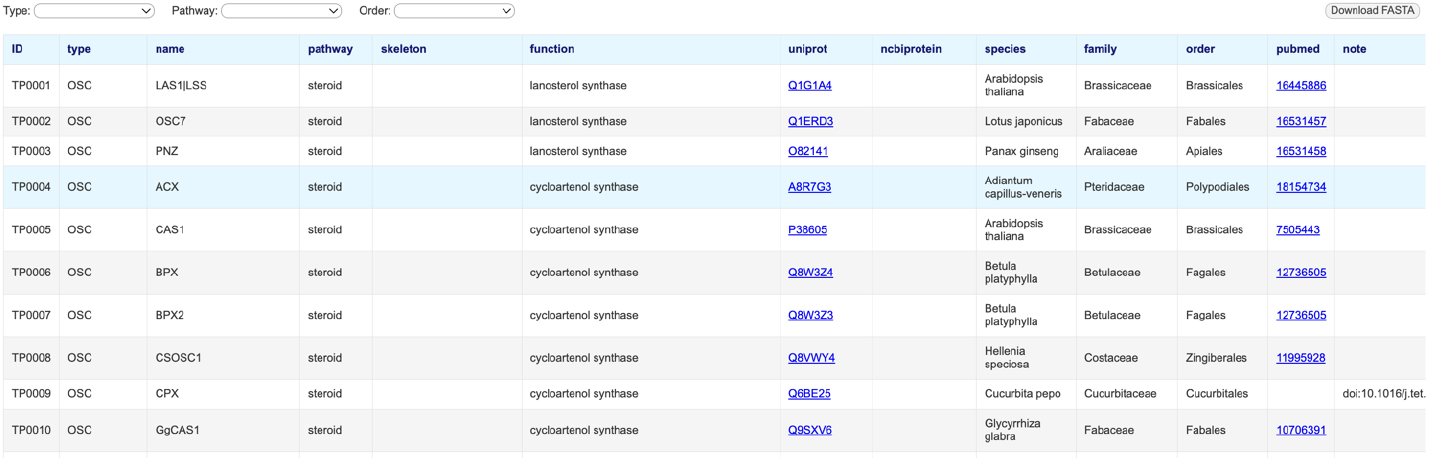
A screenshot of the database site. Drop-down menus at the top of the page work as filters for entries. Users can download amino acid sequences of the displayed data using the “Download FASTA” button.

**Table 1.**
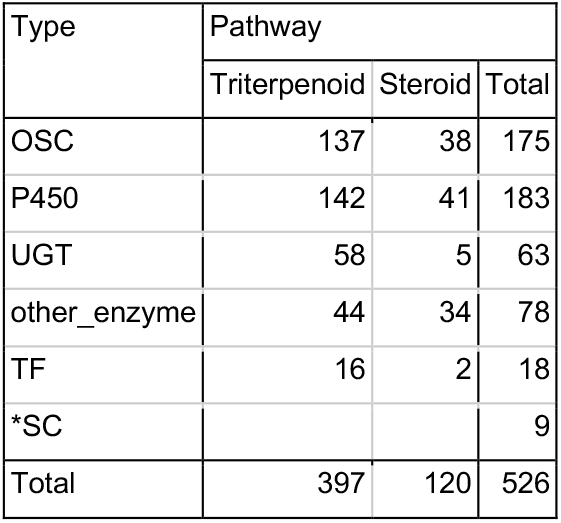
Number of protein entries in the database based on the classification of “type” and “pathway”.

**Table 2.** Number of protein entries involved in triterpenoid biosynthesis classified by taxonomic order and protein type.

| Order | Type |  |  |  |  |  |
| --- | --- | --- | --- | --- | --- | --- |
|  | OSC | P450 | UGT | other_enzyme | TF | Total |
| Apiales | 8 | 12 | 16 |  |  | 36 |
| Aquifoliales |  | 4 | 1 |  |  | 5 |
| Asterales | 21 | 6 |  | 1 |  | 28 |
| Brassicales | 16 | 11 | 10 | 5 | 7 | 49 |
| Caryophyllales | 7 | 6 | 5 | 4 |  | 22 |
| Celastrales | 6 | 7 |  |  |  | 13 |
| Cucurbitales | 10 | 9 | 1 | 2 | 2 | 24 |
| Dioscoreales |  |  |  |  |  |  |
| Ericales | 1 | 2 |  |  |  | 3 |
| Fabales | 15 | 43 | 23 | 12 | 4 | 97 |
| Fagales | 5 | 1 |  |  |  | 6 |
| Gentianales | 2 | 2 |  |  |  | 4 |
| Lamiales | 4 | 6 |  |  |  | 10 |
| Liliales |  |  |  |  |  |  |
| Malpighiales | 8 |  |  |  | 3 | 11 |
| Myrtales | 7 | 3 |  |  |  | 10 |
| Poales | 6 | 7 | 2 | 4 |  | 19 |
| Polypodiales |  |  |  |  |  |  |
| Ranunculales | 1 | 2 |  |  |  | 3 |
| Rosales | 5 | 3 |  |  |  | 8 |
| Sapindales | 6 | 13 |  | 16 |  | 35 |
| Saxifragales | 4 |  |  |  |  | 4 |
| Solanales | 4 | 3 |  |  |  | 7 |
| Vitales |  | 2 |  |  |  | 2 |
| Zingiberales | 1 |  |  |  |  | 1 |

**Table 3.** Number of protein entries involved in steroid biosynthesis classified by taxonomic order and protein type.

| Order | Type |  |  |  |  |  |
| --- | --- | --- | --- | --- | --- | --- |
|  | OSC | P450 | UGT | other_enzyme | TF | Total |
| Apiales | 2 |  |  |  |  | 2 |
| Aquifoliales |  |  |  |  |  |  |
| Asterales | 1 |  |  |  |  | 1 |
| Brassicales | 2 | 11 |  | 17 |  | 30 |
| Caryophyllales |  |  |  |  |  |  |
| Celastrales | 3 |  |  |  |  | 3 |
| Cucurbitales | 5 |  |  |  |  | 5 |
| Dioscoreales |  | 4 |  |  |  | 4 |
| Ericales |  |  |  |  |  |  |
| Fabales | 7 | 4 |  | 3 |  | 14 |
| Fagales | 2 |  |  |  |  | 2 |
| Gentianales |  |  |  |  |  |  |
| Lamiales |  | 1 |  |  |  | 1 |
| Liliales | 1 | 9 |  | 1 |  | 11 |
| Malpighiales | 5 |  |  |  |  | 5 |
| Myrtales | 2 |  |  |  |  | 2 |
| Poales | 1 | 5 |  | 1 |  | 7 |
| Polypodiales | 3 |  |  |  |  | 3 |
| Ranunculales |  | 1 |  |  |  | 1 |
| Rosales |  |  |  |  |  |  |
| Sapindales |  |  |  |  |  |  |
| Saxifragales | 1 |  |  |  |  | 1 |
| Solanales | 2 | 6 | 5 | 12 | 2 | 27 |
| Vitales |  |  |  |  |  |  |
| Zingiberales | 1 |  |  |  |  | 1 |

The future direction for Triterpene RDF is to annotate proteins using either pathway or reaction databases. For example, UniProtKB uses Rhea (http://www.rhea-db.org) (Lombardot et al. 2019) for enzyme annotation, which eases the process of integrating metabolites with protein information (Morgat et al. 2020). Such linked data would render it possible to obtain a set of annotated protein sequences necessary for the biosynthesis of a specified triterpene compound; however, the coverage of reactions for plant triterpene biosynthesis in the Rhea database is limited. One possible solution is to deposit the relevant pathways necessary for mapping the entries for this triterpene database in the community-driven pathway database WikiPathways (https://www.wikipathways.org/) (Agrawal et al. 2023). Although the current scope of WikiPathways mainly covers model organisms, efforts to expand the database to include non-model organisms are ongoing (Pico et al. 2023; Oec et al. 2023). Future integration with this type of pathway or reaction database will enforce the annotation and characterization of triterpene biosynthetic genes.

## Abbreviations

OSC: oxidosqualene cyclase
P450: cytochrome P450 monooxygenase
RDF: Resource Description Framework
SC: squalene cyclase
TF: transcription factor
UGT: UDP-dependent glycosyltransferase

## Acknowledgements

We thank the developers and organizers who attended the domestic BioHackathons in Japan (BH22.9 and BH23.9, organized by Database Center for Life Science (DBCLS)) and the Togothon meetings (organized by DBCLS) for their helpful discussions and technical support.

## Author contribution

Conceptualization: K.T.; Methodology, K.T. and H.C.; Software, K.T. and H.C.; Validation: K.T., H.C.; Formal analysis, K.T., H.C.; Investigation, K.T., H.C.; Resources, K.T., H.C., and H.B.; Data curation, K.T. and H.C.; Visualization: K.T. and H.C.; Validation: K.T. and H.C.; Resources: K.T., H.C., and H.B.; Writing—original draft: K.T.; Writing—review and editing: K.T., H.C., and H.B.; Supervision: H.B.; Project administration: K.T.; Funding acquisition: K.T. and H.B.

